# Symptom Severity in Youths with Attention Deficit Hyperactivity Disorder Associated with Normalizing Effects of Treatment on fMRI Response during a Stop Signal Task

**DOI:** 10.1101/599803

**Authors:** Catherine E. Hegarty, Mohan W. Gupta, Eric Miller, Kevin Terashima, Sandra Loo, James McCracken, James McGough, Robert M. Bilder, Russell A. Poldrack, Adriana Galvan, Susan Bookheimer

## Abstract

Inhibitory control deficits represent one of many core cognitive deficits in Attention-Deficit/Hyperactivity Disorder (ADHD). Neuroimaging studies suggest that individuals with ADHD exhibit atypical engagement of neural systems during response inhibition, but the exact nature of this phenotype is obscured by mixed findings. We tested whether drug-free youths with ADHD (n=30, ages 7-14 years, 10 female) exhibited atypical neural correlates of response inhibition, as measured with a stop signal task and fMRI, compared to matched controls. We next investigated medication effects and whether there was a relationship between symptom severity and medication effects on the fMRI-evaluated signal. Finally, we tested for a significant difference between effects of monotherapy and combined pharmacological treatment. Patients showed significantly slower stop signal response time and lower percent inhibition, but no significant differences in the neural correlates of response inhibition relative to controls. However, patients showed significantly elevated signal in frontostriatal regions during responses. Prefrontal signal in patients was positively associated with reaction time variability in patients, and change (medicated – drug free) in the prefrontal signal was significantly associated with symptom scores, such that patients with elevated symptoms had greater BOLD signal reduction following treatment. Medication significantly improved go response time median and variability as well as stop signal reaction time, but there were no significant effects of medication or treatment type on BOLD signal. These findings challenge the notion of frontostriatal hypoactivation during response inhibition as a biomarker for ADHD and suggest that symptom severity may be associated with response to medication.

## Introduction

Attention deficit hyperactivity disorder (ADHD) is one of the most common psychiatric disorders in children and adolescents, with recent global prevalence estimates reaching as high as 7.2% (Thomas et al. 2015). This challenging disorder is characterized by a pattern of debilitating impairments across a number of cognitive domains, including response inhibition. Although reduced brain activity during response inhibition has been proposed as a putative neurobiological marker for ADHD (Hart et al. 2013), neuroimaging findings for response inhibition tasks have been mixed. For example, a comparable number of studies have reported hypo- (Durston et al. 2003; Janssen et al. 2015; Konrad et al. 2006; Rubia et al. 2010) and hyperactivation (Massat et al. 2018; Pliszka et al. 2006; Schulz et al. 2004) of frontostriatal networks during response inhibition. The Stop-Signal Task (SST) (Iaboni et al. 1995) is a well-established paradigm for evaluating response inhibition that requires inhibiting a response upon presentation of an auditory cue. A 2013 meta-analysis of five fMRI studies employing the SST indicated that ADHD was associated with hypoactivation of inferior, middle and superior frontal gyri relative to a control sample (Hart et al. 2013). However, multiple studies not included in that analysis have reported contradictory results (Pliszka et al. 2006; Schulz et al. 2004).

A cohesive explanation of the neurobiology underlying aberrant response inhibition is further complicated due to clinically and developmentally heterogeneous samples. Medication status is also inconsistent across studies, with heterogeneous samples of drug-naïve, drug-free and medicated individuals. Here, we compared a cohort of 30 drug-free youth, with both inattentive and combined type ADHD, to a matched sample of typically developing youth. We used the SST paradigm in conjunction with fMRI to evaluate differences in SST performance and neural correlates of response inhibition in youths with ADHD and healthy controls. We then investigated medication effects on the task in patients and determined whether any such effects were associated with symptom severity. Finally, we explored whether there were any significant differences between mono-and combined-pharmacotherapy.

## Methods

### Participants

Participants were recruited through UCLA’s Translational Research to Enhance Cognitive Control (TRECC) research center. Parents and participants provided written informed permission and assent. All study procedures were approved by the UCLA Institutional Review Board and overseen by a Data Safety and Monitoring Board. Details of inclusion criteria can be found in Supplemental Data. Details of the 8-week randomized, double-blind randomized controlled trial procedures have been previously described (McCracken et al. 2016). Briefly, youths with ADHD were assigned to treatment with either d-methylphenidate (DMPH), guanfacine (GUAN) or combined treatment (COMB) with both DMPH and GUAN. Final assessments were eight weeks after the initial baseline visit. A subset of youths enrolled in the larger study (McCracken et al. 2016) performed the SST while undergoing fMRI scanning (n=51 controls and n=106 ADHD). Youths with ADHD were only included in this analysis if they successfully completed both baseline and follow-up scans. After assessing for additional exclusion criteria (see Supplemental Data), a final sample of 30 youths with ADHD (age =10.12 ± 1.69 years; 10 female) remained for analysis. Treatment and ADHD-subtype distributions are listed in Table 1. A control sample of 30 individuals was selected from the usable control group to match the ADHD sample on age and sex (Table 1). Neuropsychological data was also acquired at baseline visits, including the scores for Strengths and Weakness of ADHD Symptoms and Normal Behavior Rating (SWAN).

**Table 1.** Demographic information

|  | **ADHD**<br>**(n=30)** | **CTL**<br>**(n=30)** | **Statistic** | **Significance** |
| --- | --- | --- | --- | --- |
| Age±SD(years) | 10.12±1.69 | 10.23±1.77 | t(58)=-0.254 | p=0.800 |
| Female (%) | 10 (33.3%) | 10 (33.3%) |  |  |
| Race | | | $\chi^2(3, N=60)=1.026$ | p=0.795 |
| Caucasian | 20 | 19 |  |  |
| African American | 7 | 7 |  |  |
| Asian | 0 | 1 |  |  |
| Other | 3 | 3 |  |  |
| Ethnicity | | | $\chi^2(1, N=60)=1.18$ | p=0.278 |
| Hispanic or Latino | 3 | 6 |  |  |
| Not Hispanic or Latino | 27 | 24 |  |  |
| SWAN±SD<sup>1</sup> | 65.8±17.9 | 115.3±36.7 | t(55)=-6.56 | p=1.93x10<sup>-8</sup> |
| ADHD Subtype (%) |  |  |  |  |
| Combined | 17 (56.7%) |  |  |  |
| Inattentive | 13 (43.3%) |  |  |  |
| Medication (%) |  |  |  |  |
| DMPH | 13 (43.3%) |  |  |  |
| GUAN | 7 (23.3%) |  |  |  |
| DMPH+GUAN | 10 (33.3%) |  |  |  |

<sup>1</sup>SWAN data was only available for 27 of the 30 control participants
GUAN=guanfacine, DMPH= d-methylphenidate, SWAN= Strengths and Weakness of ADHD Symptoms and Normal Behavior Rating

### Behavioral *Analysis*

Additional details of the SST employed and behavioral inclusion criteria can be found in Supplemental Data. Independent t-tests were conducted to test for group differences on median reaction time (RT), RT standard deviation (used as a measure of RT variability [RTV]), percentage of successful inhibition and accurate Go trials, and stop signal reaction time (SSRT). Paired t-tests were used to evaluate medication effects on these behavioral measures within the ADHD group.

### Image Acquisition and Preprocessing

See Supplemental Data for details of image acquisition and preprocessing.

### Contrasts of Interest

Go, Successful Stopping (SuccStop), Unsuccessful Stopping (UnsuccStop), and nuisance events (missing responses or errors on Go trials) were modeled after convolution with a canonical hemodynamic response function (HRF). For each participant, three principle contrasts of interest were computed comparing each event — Go, SuccStop and UnsuccStop — to implicit baseline. Significant between-group differences in any of these principle contrasts were then used to mask additional contrasts involving the event of interest. For example, significant group differences for Go > Baseline were used as a pre-threshold mask for the contrasts of Go > Succstop and Go > UnsuccStop. In the absence of significant between-group findings for one of the three principle contrasts (e.g. Event X > baseline), no further post-hoc analyses (e.g. Event X > Event Y or Event X > Event Z) relating to that contrast were run.

### Statistical Analyses

Mixed effects analyses in FSL’s FEAT were used to combine runs within subjects for individuals with two usable runs. These mid-level analyses and first-level analyses (from individuals with only one usable SST run) were fed into higher-level analyses. Ten individuals with ADHD and nine individuals from the control group only had one usable SST run; the total number of runs included did not significantly differ between groups (t(60)=-0.273, p=0.786). Group analyses were conducted using a general linear model in FSL’s FLAME1 with a significance threshold of Z > 3.0. Peak coordinates of significant clusters in between-group comparisons were used to generate regions of interest for post hoc analyses (described below). An insufficient number of the matched control group (n=13) returned for follow-up visits, so baseline data from control participants were compared to the ADHD group at both baseline and follow-up visits. Overall effects of medication (all treatment types combined) within the ADHD group were evaluated using a paired t-test in FSL’s FEAT.

### Post-hoc Analyses

Difference images (follow-up minus baseline) for each contrast of interest were created using AFNI’s 3dcalc. For contrasts with significant between-group results, average time-series from 5mm spherical regions of interest (ROI) around peak voxels were extracted from the baseline and difference images. To further explore effects of treatment type of change in ROI signal between visits, we used a multivariate analysis with the extracted difference score (Δ_BOLD_) at each ROI as the dependent variable, treatment type (GUAN, DMPH, COMB) as the fixed factor and age, sex and days between visits as covariates. Next, partial correlations, controlling for age, sex, and days between visits, were run to evaluate whether baseline symptom severity (i.e. SWAN score) was associated with response to treatment (Δ_BOLD_). Spearman’s rank correlations were used to account for non-normality of the data. Finally, given the importance of RTV in response inhibition abnormalities associated with ADHD (Tamm et al. 2012), we evaluated the relationship between RTV and the extracted baseline ROI values for both patients and controls. All statistical tests were conducted in SPSS (v. 25) with a significance threshold of p<0.05.

## Results

Youths with ADHD had significantly higher SSRT (t(58)=2.37, p=0.021) and lower percent inhibition (t(58)=-2.83), p=0.006) than matched controls. There was also a trend-level difference in RTV (t(58)=-1.77, p=0.083). However, there were no significant differences between groups in median RT or the average percent of go trials answered (Table S1).

Youths with ADHD showed significantly higher signal for the Go > Baseline contrast than controls in the left middle frontal gyrus (MFG), right superior frontal gyrus (SFG), and left putamen (Figure 1, Table S2). However, there were no significant differences between controls and the ADHD group at follow-up. There were no significant between group differences at either baseline or follow-up for any other contrast, nor were there any significant differences between drug-free and medicated conditions within the ADHD group.

**Figure 1.**
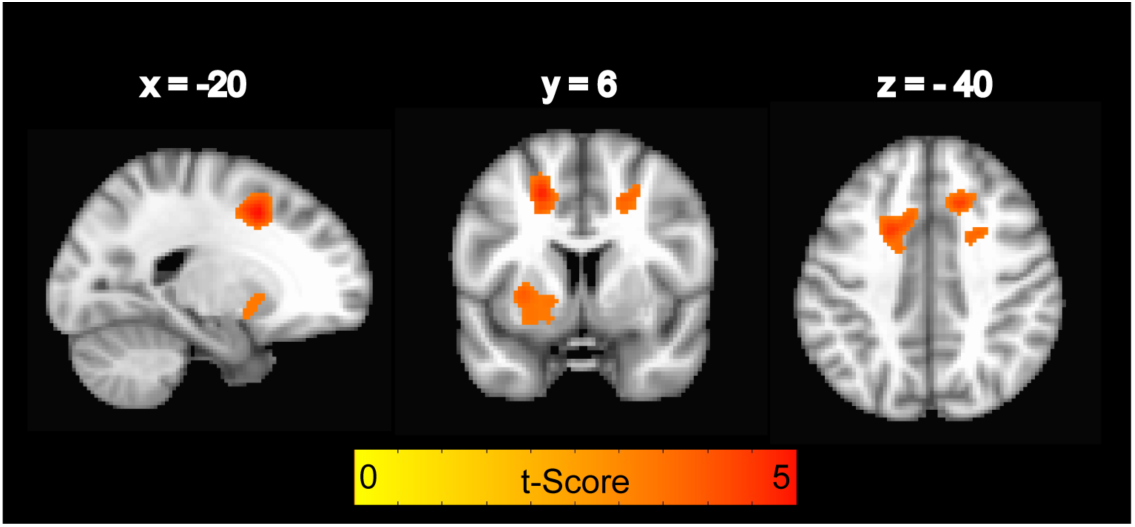
Voxelwise between group results: ADHD > Control for Go > Baseline contrast. Image is thresholded at p<0.001.

Partial correlations, controlling for age and sex, indicated that baseline RTV was significantly associated with baseline MFG signal (r_s_(n=30)=-0.620, p=4.3×10^-4^; Figure 2A) and had a trend-level association with baseline SFG signal (r_s_(n=30)=-0.344, p=0.073; Figure 2B). The control group did not show any such relationship (p>0.10).

**Figure 2.**
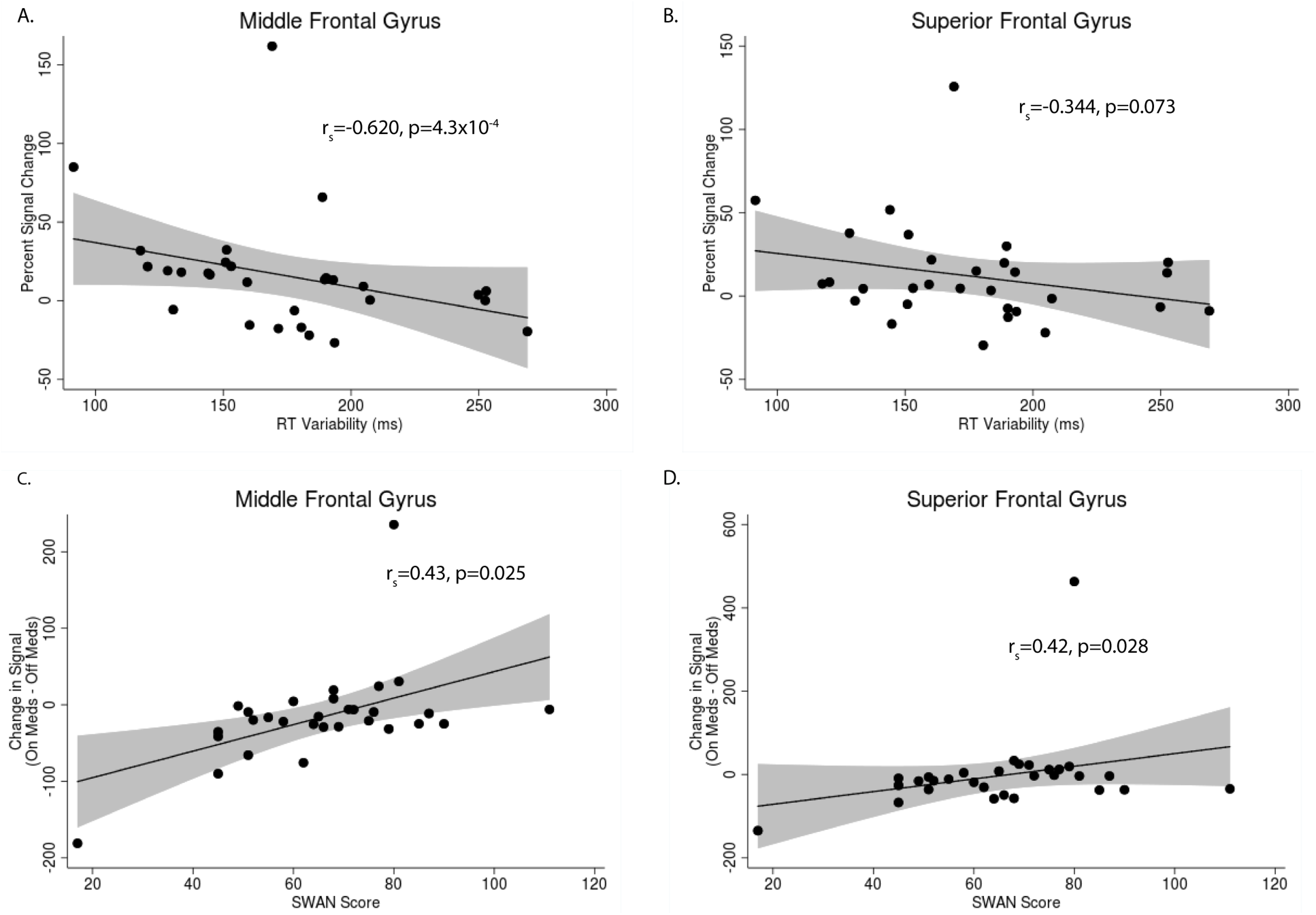
Inverse relationships in ADHD between reaction time (RT) variability and percent signal change in the (A) left MFG and (B) right SFG for the Go > Baseline contrast. Signal Changes (Medicated – Drug Free) in the (C) left MFG and (D) right SFG show a positive association with baseline SWAN, such that more severe symptoms were associated with a signal decrease after medication.

Medication was associated with significantly faster median RT (t(29)=3.41, p=0.002), less RTV (t(29)=3.67, p=0.001), and shorter SSRT (t(29)=4.59, p=7.9×10^-5^) relative to drug-free performance at baseline (Table S4). There were also significant correlations between SWAN scores and Δ_BOLD_ for the left MFG (r_s_(n=30)=0.429, p=0.025; Figure 2C) and right SFG (r_s_(n=30)=0.422, p=0.028; Figure 2D), such that lower SWAN scores were associated with a greater reduction in signal following treatment. There was no significant relationship between SWAN score and Δ_BOLD_ in the left putamen. There were no significant effects of treatment type (DMPH vs. GUAN vs. COMB) on any changes in performance or Δ_BOLD_ for any of the ROIs.

## Discussion

Compared to a matched sample of typically developing youths, drug-free youths with ADHD show significantly elevated BOLD signal during the Go > Baseline condition in frontostriatal regions previously implicated in response inhibition (Verbruggen and Logan 2008). Although there are limited studies exploring the Go > Baseline contrast with which to compare the findings, abnormal motor activity (assessed with cardiac response patterns) has been previously reported in youths with ADHD during the “go” condition of a go/no-go task (Borger and van der Meere 2000). We did not find any significant differences in the neural correlates of contrasts involving inhibition; however, our findings show that youths with ADHD have significantly slower SSRTs and lower percent inhibition. This apparent divergence of the imaging and behavioral data has also been found in previous studies. For example, five of the studies reporting abnormal neural correlates of response inhibition found either no corresponding deficits in SST performance (Cubillo et al. 2010; Pliszka et al. 2006; Rubia et al. 2010) or reduced probability of inhibition without differences in SSRT (Rubia et al. 2011; Rubia et al. 1999). However, our finding of an association between RTV and abnormal prefrontal signal during the Go condition in patients supports existing dialogue about the role of RTV as a more meaningful index of ADHD pathology than standard behavioral response inhibition measures.

Our finding of diminished performance without corresponding atypical brain activation may also be due to the adaptive nature of this SST design or the higher proportion of inattentive-type patients included in this sample. Of the existing neuroimaging studies using SST to investigate childhood ADHD, only one study reported inclusion of inattentive-type (11% of the total sample (Janssen et al. 2015)), while the rest exclusively included combined-type patients (Massat et al. 2018; Passarotti et al. 2010; Pliszka et al. 2006; Rubia et al. 2011). Our collective behavioral and neuroimaging findings may thus support existing theories that slower SSRT does not reflect poor motor inhibition, but is rather a reflection of deficits in attention and cognitive processing driven by the inattentive-type patients (Alderson et al. 2007; Lijffijt et al. 2005), while diminished brain activity reported in other studies may be predominantly driven by the more severe hyperactive symptoms present in combined-type patients. Greater RTV has also been more closely associated with the combined than inattentive subtype (for review see (Tamm et al. 2012)), which may explain why we found only trend-level differences in RTV at the group level. Furthermore, the differences in neural correlates of response inhibition between control and ADHD youths are thought to increase with age (Hart et al. 2013), and thus it’s possible that our sample— with over half of the individuals under the age of 10 years (n=17)— was too young to reveal significant group differences.

Meta-analytic results of response inhibition reveal that effect size differences in children are quite small across most regions (Hart et al. 2013). Such modest effects, even without two additional discrepant studies (Pliszka et al. 2006; Schulz et al. 2004), calls into question the usual interpretation that frontostriatal hypoactivation during response inhibition is a hallmark deficit of ADHD and suggests possible unidentified confounders. One possibility is the high rate of ADHD comorbidity with other disorders, particularly a comorbidity with Conduct Disorder. For example, earlier SST studies reporting frontostriatal hypoactiviation included ADHD children with rates of Conduct Disorder as high as 31 – 43% (Rubia et al. 1999; Rubia et al. 2005).

Although medication improved SSRT, RT and RTV, there was no significant overall effect of medication on neural correlates of response inhibition. However, the degree to which prefrontal signal during Go > Baseline normalized (i.e. decreased) following treatment was significantly associated with the severity of symptoms at baseline: individuals with elevated symptoms showed a larger effect of treatment. The absence of any overall medication effects on BOLD signal may be due to our inclusion of individuals treated with guanfacine, which employs a post-synaptic α_2A_-agonist mechanism to increase norepinephrine rather than increasing endogenous dopamine, as is done with stimulants. While there were no significant effects of treatment type on any of our response inhibition metrics, our ability to evaluate treatment type was somewhat limited by our small guanfacine sample. However, our finding is consistent with a previous report (using the larger non-imaging data from this study) which reported no effect of treatment type on response inhibition (Bilder et al. 2016).

There are several limitations worth noting. First, our exclusion criteria excluded a large number of subjects and thus results from the remaining sample may not be representative of the larger group. Second, due to the nature of the clinical trial, patients were not counterbalanced for treatment order and thus were all drug-free at baseline and medicated at the follow-up visit. Therefore, we cannot rule out any potentially confounding effects of repeated testing. Third, we did not have sufficient data at the follow-up visit for the control group to compare changes between baseline and follow-up to the ADHD group. Although we did not find any significant differences between the ADHD group at baseline (drug-free) and follow-up (medicated), the absence of any significant difference between the control group at baseline and the ADHD group at follow-up may be due to the fact that it was the second time the ADHD group had undergone SST in the scanner. Last, our evaluation of treatment type effects was limited by a disproportionately small sample in the guanfacine group. Future studies with larger treatment groups would be better powered to investigate these effects.

Collectively, these results challenge the belief that hypoactivation of the frontostriatal network during response inhibition on the SST is a biomarker of ADHD. Furthermore, we provide evidence that baseline symptom severity is a predictor of response to treatment with DMPH and/or GUAN, regardless of whether it is combined or monotherapy.

