## Supplementary Data for "Symptom Severity in Youths with Attention Deficit Hyperactivity Disorder Associated with Normalizing Effects of Treatment on fMRI Response during a Stop Signal Task"

*Inclusion and Exclusion criteria*

Inclusion criteria for the ADHD group were 1) male or female aged 7-14 years; 2) DSM-IV ADHD (any subtype) diagnosed by the Kiddie-Schedule for Affective Disorders and Schizophrenia-PL and clinical interview; and 3) Clinical Global Impression—Severity (CGI-S) score ≥ 4 for ADHD. Exclusion criteria for participants were 1) autistic disorder, chronic tic disorder, psychosis, bipolar disorder, or structural heart defects; 2) current major depression or panic disorder; 3) systolic or diastolic blood pressure > 95th or <5th percentile for age and body mass index (BMI); 4) medical condition contraindicating stimulants or alpha agonists; 5) need for chronic use of other CNS medications. A healthy comparison group was comprised of boys and girls aged 7-14 who had no lifetime history of DSM-IV-TR Axis I mental disorders as determined by KSADS-PL and clinical interview, and who had exclusion criteria noted above.

Youths with ADHD were only included in this analysis if they successfully completed both baseline and follow-up scans at 8-weeks after the baseline visit. To ensure that only the highest quality data were included in analyses, a subject’s neuroimaging data was excluded from analysis if there was excessive (>3.5mm) relative head motion, as estimated by FSL’s MCFLIRT motion correction during preprocessing of the raw data, at any time point during both runs (baseline: ADHD n=25; CTL n=3). Remaining subjects were also excluded if they did not meet the minimum behavioral performance threshold (baseline: ADHD n=13; CTL n=2; see Behavioral Analysis section for exclusion criteria). Finally, ADHD participants were only selected for analysis if they had usable data for both baseline and follow-up visits. Of the 68 patients with usable baseline data, 41 also had follow-up visit data. Of those 41, eleven were excluded for excessive head motion. The remaining 30 patients with data at both baseline and follow-up visits all met behavioral performance criteria and were thus included in analyses.

*Stop Signal Task*

During scanning, participants performed the SST (Figure S1). On each trial, a left or right pointing arrow appeared on the screen. Participants were instructed to respond as quickly as possible with a left or right key press. On 25% of trials, a stop signal (auditory tone, 900 Hz, 500 milliseconds [ms] duration) was sounded, signaling the participant to withhold his/her response. After the response or 1 sec (if inhibition was successful), the stimulus disappeared and was followed by a jittered delay (0.5 to 4 s, mean = 1s). The interval between the stimulus and the stop signal (i.e., stop signal delay, SSD) was varied with each participant’s performance. Average stop signal duration (SSD) was computed for each participant from the values of the two staircases after each converged on 50% inhibition. The SSD was increased 50 ms after successful inhibition, and decreased 50 ms after inhibition failure. This procedure ensured that subjects successfully inhibited responses on approximately 50% of inhibition trials, which allows computation of the stop signal reaction time and ensures that both behavioral performance and proportion of successful stop trials were equated across participants.

The initial SSD was selected from one of two interleaved algorithms (staircases), each starting with SSD values of 200 and 320 ms and then increased or decreased according to the participant’s performance. In Run 2, the last SSD of each staircase on the first run was used as the starting value of each respective staircase. There were 96 Go and 32 Stop trials per run, with equal numbers of leftward and rightward-pointing arrows. Participants performed two runs of the task (256 trials total).

For subjects with two usable runs, SSRT was averaged across both runs. For a run to be included in the analysis, participants were required to respond on greater than 60% of Go trials, have greater than 90% accuracy on Go trials, and 25-75% successful inhibition. Any responses under 50 ms were not included in the analyses, which is common practice in response inhibition literature (Congdon et al. 2012).

Events (1 sec) were modeled at stimulus onset. Null events, consisting of the jittered inter-trial interval when the screen was blank, were not explicitly modeled and therefore constituted an implicit baseline.

*Image Acquisition*

Data were acquired on a 3T Siemens Trio MRI scanner. For each run, 182 functional T2*-weighted echoplanar images (EPI) were acquired [slice thickness, 4 mm; 34 slices; TR, 2 sec; TE, 30 mec; flip angle, 90°; matrix, 64 x 64; FOV, 200 mm; voxel size, 3 x 3 x 4 mm3]. Two volumes, collected at the beginning of each run to allow for T1 equilibrium effects, were discarded. A T2-weighted, matched-bandwidth (MBW), high-resolution, anatomical scan and magnetization-prepared rapid-acquisition gradient echo (MPRAGE) were acquired for each subject for registration (TR, 2.3; TE, 2.1; FOV, 256; matrix, 192 x 192; sagittal plane; slice thickness, 1 mm; 160 slices). The orientation for MBW and EPI scans was oblique axial to maximize brain coverage. Matlab and Psychtoolbox ([www.psychtoolbox.org](http://www.psychtoolbox.org/)) were used for stimulus presentation and timing.

*Image Preprocessing*

All fMRI preprocessing and analyses were performed using FSL version 5.0.9 (http:www.fmrib.ox.ac.uk/fsl). Images were skull-stripped using FSL’s brain extraction tool (BET) and spatially smoothed using an 8 mm FWHM Gaussian kernel. FSL’s MCFLIRT was used for motion correction. A high pass filter cut off of 90 seconds was applied. Three-step registration was performed using the FLIRT software to register the EPI images to the matched-bandwidth structural image, then the MPRAGE structural image, and finally the standard MNI152 space using a 12-degree of freedom (DOF) affine transformation, with a resampling resolution of 4mm. FSL’s Multivariate Exploratory Linear Decomposition into Independent Components (MELODIC version 3.15) was used to run single-session independent component analysis (ICA) on each SST run and estimate spatial ICAs by maximizing non-Gaussian sources with robust voxelwise variance normalization of time courses, automatic dimensionality estimation, and an IC map threshold of 0.5. Independent components for runs from 10 patients and 10 controls were manually labeled as either signal or noise (e.g. susceptibility and motion artifacts, cardiac contributions) according to predefined criteria (Griffanti et al. 2017) by two independent raters (CH and MG), with greater than 90% agreement. These 20 manually-labeled runs were used as a training dataset for FMRIB’s ICA-based Xnoiseifier (FIX version 1.065; MATLAB v. 8.6; R 3.4.0) to clean the remaining usable runs (Griffanti et al. 2014; Salimi-Khorshidi et al. 2014). To maximize the amount of signal included, a threshold of 5 (mean true positive rate = 96.1%) was applied. FIX-cleaned images were then used for subsequent lower-level statistical processing.

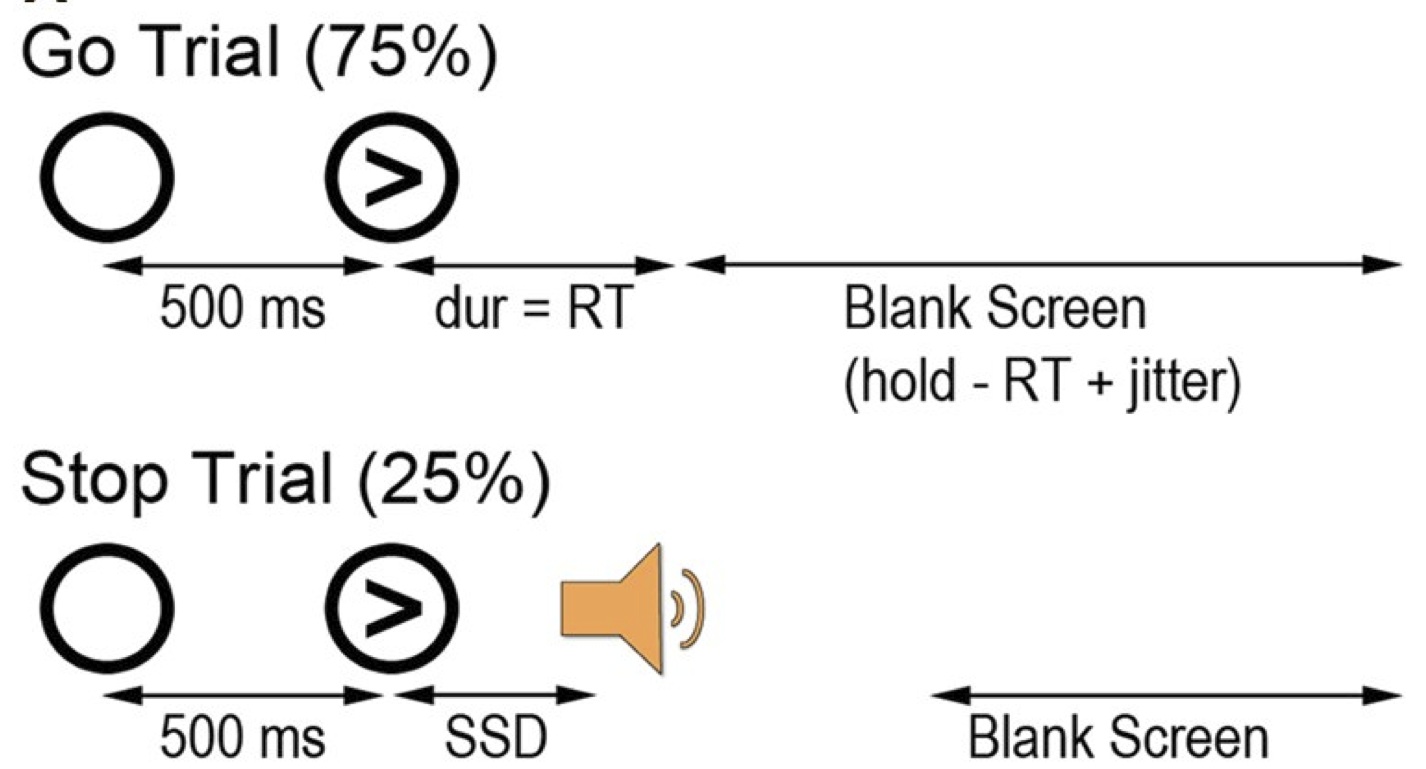
Figure S1. Schematic of Stop-signal Task

.

Table S1. Behavioral Results

| **Measure** | **ADHD (n=30)** | **CTL (n=30)** | **Statistic** | **Significance** |
| --- | --- | --- | --- | --- |
| SSRT ± SD (ms) | 299.3 ±79.2 | 247.9±88.3 | t(58)=2.37 | p=0.021 |
| Median RT± SD (ms) | 566.2±108.3 | 583.0±102.7 | t(58)=-0.617 | p=0.540 |
| RT Variability ± SD (ms) | 175.0±42.8 | 157.5±33.4 | t(58)=1.77 | p=0.083 |
| % Inhibition± SD | 49.1±6.2 | 54.0±7.1 | t(58)=-2.83 | p=0.006 |
| % Go Answered± SD | 95.4±5.5 | 97.1±3.8 | t(58)=-1.36 | p=0.180 |

RT=Reaction time, SD=standard deviation, SSRT=stop signal reaction time

Table S2. Voxelwise Results for Contrasts of Interest

| **Brain Region** | **Voxel** | **x** | **y** | **z** | **Z** |
| --- | --- | --- | --- | --- | --- |
| **Go > Baseline** | | | | | |
| **ADHD > Controls** | | | | | |
| L MFG | 790 | -20 | 10 | 44 | 5 |
| L Putamen | 698 | -26 | 16 | 4 | 4.01 |
| R SFG | 469 | 14 | 24 | 40 | 4.41 |
| **Controls** | | | | | |
| L Precentral Gyrus | 1476 | -38 | -22 | 64 | 5.22 |
| **ADHD** | | | | | |
| L Precentral Gyrus | 5313 | -40 | -22 | 56 | 7.59 |
| R Cerebellum | 1110 | 22 | -50 | -24 | 6.05 |
| L Putamen | 840 | -24 | 4 | 0 | 5.62 |
| R LOC | 513 | 48 | -70 | -4 | 5.0 |
| **Successful Stop > Baseline** | | | | | |
| **Controls** | | | | | |
| L Insula | 2331 | -36 | 14 | 8 | 5.7 |
| R IFG (pars Opercularis) | 1738 | 48 | 6 | 4 | 5.73 |
| R Paracingulate Gyrus | 1016 | 10 | 12 | 46 | 5.2 |
| R STG | 805 | 68 | -28 | 10 | 5.02 |
| **ADHD** | | | | | |
| L Insula | 11360 | -36 | 14 | 10 | 6.54 |
| R STG | 8561 | 70 | -30 | 14 | 6.03 |
| **Unsuccessful Stop > Baseline** | | | | | |
| **Controls** | | | | | |
| R Insula | 1275 | 36 | 14 | 4 | 5.41 |
| L ACC | 1067 | -10 | 8 | 38 | 4.46 |
| L Insula | 961 | -30 | 20 | 10 | 4.98 |
| L Postcentral Gyrus | 559 | -44 | -18 | 64 | 4.3 |
| **ADHD** | | | | | |
| L Insula | 4974 | -34 | 12 | 10 | 6.68 |
| R ACC | 3910 | 8 | 12 | 38 | 6.12 |
| R Precentral Gyrus | 2528 | 52 | 8 | 4 | 5.39 |

Abbreviations: MFG= middle frontal gyrus, SFG = superior frontal gyrus, LOC = lateral occipital cortex, ACC = anterior cingulate cortex, STG = superior temporal gyrus, IFG = inferior frontal gyrus

Table S3. Medication Effects on SST Performance

| **Measure** | **Mean Difference (Drug Free - Medicated)** | **Statistic** | **Significance** |
| --- | --- | --- | --- |
| SSRT ± SD (ms) | 71.2 ± 85.0 | t(29)=4.59 | p=7.9x10^-5^ |
| Median RT± SD (ms) | 48.1±77.4 | t(29)=3.41 | p=0.002 |
| RT Variability ± SD (ms) | 29.61±44.2 | t(29)=3.67 | p=0.001 |
| % Inhibition± SD | -1.81±7.4 | t(29)=-1.34 | p=0.190 |
| % Go Answered± SD | -1.9±6.2 | t(29)=-1.71 | p=0.099 |

RT=Reaction time, SD=standard deviation, SSRT=stop signal reaction time
